# Single-Sample Network Topology Unveils Pathogenic Hubs in Sjögren’s Disease

**DOI:** 10.64898/2026.09.16.752113

**Authors:** João V. F. Cavalcante, Rodrigo J. S. Dalmolin, Diego Marques-Coelho

## Abstract

Sjögren’s disease (SD) is a systemic autoimmune condition characterized by extensive clinical and biological heterogeneity, complicating the development of targeted therapies. To better describe the personalized molecular rewiring driving SD pathogenesis, we utilized bulk RNA-sequencing data from whole blood samples of the PRECISESADS IMI consortium to construct sample-specific gene interaction networks by applying the LIONESS algorithm integrated with experimental protein-protein interaction evidence. Comparing SD patients against healthy controls, we identified an absolute total of 647 Differentially Interacting Genes (DIGs) representing structural shifts in network connectivity. Topological clustering of these DIGs revealed four major functional macro-modules underlying disease pathology: defense response to virus, B cell activation, DNA replication, and positive regulation of protein catabolic process. Central topological hubs, such as ISG15, OASL, UBE2L6, and LGALS3BP, were identified as drivers of this reorganization, a fact supported by integrating network topology with single-sample Gene Set Enrichment Analysis (ssGSEA), which demonstrated that the interaction degree of these hubs exhibits strong positive correlations with the functional activity of their respective pathogenic pathways. To translate these structural findings into therapeutic opportunities, we performed an in-silico network vulnerability analysis on patient-specific interaction networks, ranking targets by the fractional loss of global efficiency following their removal and correcting for node degree. This approach prioritised the receptor tyrosine kinase *EPHB2*, which has no prior description in this disease, alongside the kinase *BLK* and the B-cell co-receptor *CD22*, a target independently evaluated in a previous clinical trial in Sjögren’s disease. Thus, our single-sample network framework provides a computational mechanism to uncover pathogenesis-related modules and to prioritise drug repurposing candidates in SD, while also recovering targets whose clinical failure bounds what network topology alone can predict.

**Highlights:**

- Sample-specific networks reveal interactome rewiring in Sjögren’s disease.
- Hub connectivity strongly correlates with interferon pathway activity.
- Network vulnerability analysis prioritizes novel structurally critical drug targets.

## 1. Introduction

Sjögren’s Disease (SD) is a chronic, systemic autoimmune disorder primarily characterized by the infiltration of lymphocytes into the salivary and lacrimal glands, resulting in profound oral and ocular mucosal dryness Mavragani and Moutsopoulos (2020). Beyond these hallmark symptoms, the disease is defined by profound B cell hyperactivity and can involve systemic complications affecting the respiratory, neurological, and renal systems Verstappen et al. (2021). A major challenge in managing SD, as with other systemic autoimmune diseases, is its significant clinical and biological heterogeneity Barturen et al. (2021); Zhao et al. (2024). The therapeutic landscape has shifted substantially: telitacicept became the first agent approved for this indication Xu et al. (2024), and ianalumab, nipocalimab, dazodalibep and iscalimab have all reported positive controlled trials Bowman et al. (2022); Noaiseh et al. (2025); St Clair et al. (2024); Fisher et al. (2024). Yet the effect sizes remain modest, and patient-reported symptom burden has proved almost universally refractory: a network meta-analysis of 27 randomised trials comprising 2261 patients found no agent superior to placebo on the EULAR Sjögren’s Syndrome Patient Reported Index Yang et al. (2026). Consequently, there is an urgent need for more refined, personalised strategies that account for the unique molecular landscape of each patient.

To address this complexity, researchers are increasingly moving away from population-level aggregate data toward modeling biological systems at the level of the individual patient Chen et al. (2023); Deschildre et al. (2024). By extracting inherent biological information from genome-wide expression profiles, these individual-level networks characterize unique interaction patterns that reflect the specific phenotypic alterations of a single subject Kuijjer et al. (2019b). These methodologies allow for a more holistic view of biological systems and have been used successfully in very distinct biological conditions, such as to find novel subtypes in lung adenocarcinoma López-Sánchez et al. (2025) and to describe dose-dependent radiation effects in spaceflight Zhang et al. (2024).

In the context of SD, sample-specific interaction graphs provide a powerful mathematical window into the active rewiring of core pathways, such as chronic type-I and type-II interferon (IFN) cascades or downstream lymphocytic responses. By quantifying how genes change their local interaction partners between patients and controls, we can isolate specific nodes whose altered connectivity directly reflects the functional intensity or suppression of broad pathological cascades. This individualized approach moves the analytical focus from a static census of upregulated transcripts to a dynamic map of functional interaction shifts. Furthermore, exposing these individualized topology-function dependencies provides a robust computational framework for target prioritization and drug repurposing, allowing for the matching of small molecules and biologics to targets positioned close to critical, rewired cellular hubs Koutsandreas et al. (2025).

In this study, we employed a sample-specific network layout to characterize the global landscape of altered gene interactivity in SD patients compared to healthy controls. We integrated individual node connectivity with functional activity scoring to systematically uncover genes acting as structural and dynamic anchors of disease pathology, ultimately charting a rewired architectural map for downstream therapeutic target discovery.

## 2. Methods

### 2.1. Data Acquisition

The present study used bulk RNAseq data produced from whole blood samples by the PRECISESADS IMI consortium Laigle et al. (2018), comprehending two cohorts: The cross-sectional and inception cohorts. These data include both SD-diagnosed patients as well as healthy controls. SD diagnosis was performed on patients according to the 2002 American-European Consensus Group classification criteria. A detailed overview of the study’s participants can be found in Table 1.

**Table 1.** Demographic characteristics of the study cohorts.

| Cohort | Group | N | Median Age (SD) | Sex (F/M) |
| --- | --- | --- | --- | --- |
| Cross-Sectional | Healthy Control | 267 | 46 (13.0) | 202 / 65 |
|  | Sjögren's Disease | 190 | 59 (12.9) | 178 / 12 |
| Inception | Sjögren's Disease | 60 | 50 (13.2) | 57 / 3 |

### 2.2. Construction of single-sample networks

Sample-specific networks were reconstructed using the LIONESS (Linear Interpolation to Obtain Network Estimates for Single Samples) algorithm via the lionessR frame-work Kuijjer et al. (2019a). The analysis was performed using the same VST-normalized count matrix obtained before, but was restricted to only protein-coding genes. To ensure biological relevance and reduce computational noise, the resulting networks were filtered to include only edges with experimental evidence documented in the STRING database (v12.0) Szklarczyk et al. (2023). A Gene-by-Sample Degree Matrix was then constructed by applying a sample-specific threshold (mean edge weight + 2 standard deviations); for each sample, the “degree” of a gene was defined as the number of its connections exceeding this threshold.

### 2.3. Identification of Differentially Interacting Genes

In a methodology originally described by Zhang et al. (2024), in order to identify genes with significant shifts in network connectivity, we performed a Differentially Interacting Gene (DIG) analysis on the degree matrix, comparing SD patients versus healthy controls. Statistical significance was assessed using two-tailed t-tests. For each gene, Levene’s test was first applied to evaluate the homogeneity of variance; Welch’s t-test was utilized if the assumption of equal variance was violated, while Student’s t-test was used otherwise. P-values were adjusted for multiple testing using the Benjamini-Hochberg (BH) method. Genes with an adjusted p-value (*p*_*adj*_ < 0.05) were defined as DIGs.

### 2.4. Single-sample gene set enrichment analysis

Functional activity at the single-sample level was estimated using single-sample Gene Set Enrichment Analysis (ssGSEA) as implemented in the GSVA R package Hänzelmann et al. (2013). We utilized the Hallmark gene sets from the Molecular Signatures Database (MSigDB v2026.1) Subramanian et al. (2005); Liberzon et al. (2015). To identify pathways associated with SD, we applied linear modeling via the limma package Ritchie et al. (2015). The SD samples were compared against the controls to identify pathways with significantly differential enrichment scores (*p*_*adj*_ < 0.05).

### 2.5. Gene-to-Pathway Correlations

To integrate network topology with biological function, we calculated the correlation between gene connectivity (degree) and pathway activity (ssGSEA scores). Significant DIGs were correlated with the activity scores of significantly enriched Hallmark pathways using Spearman’s rank correlation. This approach identified genes whose network importance directly scales with the activation or repression of specific pathological biological processes.

### 2.6. Network Vulnerability Analysis and Drug Target Mining

To translate topological alterations into pharmacological hypotheses, we conducted an in-silico network vulnerability analysis on the collapsed DIG protein-protein interaction graph. Because edge confidence scores are similarity measures whereas shortest-path algorithms interpret edge weights as distances, we weighted the graph using the LIONESS single-sample edge weights themselves. Edge weights were recomputed for the graph edges using the estimator described above, and the median across SD patients and, separately, across healthy controls was converted to a distance as *d* _*ij*_ = 1 − |*w* |_*ij*_.

Vulnerability was quantified as the fractional decrease in the network’s Global Efficiency (GE), defined as the average inverse shortest path length between all node pairs Fan et al. (2022), following the deletion of each candidate node. Because raw deletion scores nonetheless retain a correlation with node degree, we additionally report the residual of a linear regression of vulnerability on degree, which identifies nodes contributing more than their connectivity alone predicts.

Candidate targets were DIGs present in the PPI graph for which DrugBank version 5.1.13 Wishart et al. (2018) lists at least one approved drug with an annotated action on the target (e.g. inhibitor, antagonist, agonist). Ensembl gene IDs were mapped to UniProt accessions using the org.Hs.eg.db package Carlson. Annotations to non-specific compounds, such as metal ions, metabolic cofactors and antibodies linked to the target only through Fc binding, were disregarded. Y-linked and X-inactivation-escape genes were also excluded, as their connectivity reflects the different sex ratios of the patient and control groups rather than disease. Target annotations from ChEMBL release 36 Zdrazil et al. (2024) were additionally consulted when selecting representative drugs.

## 3. Results

To investigate the systemic reorganization of the gene interaction architecture in SD, we employed the LIONESS algorithm to reconstruct sample-specific co-expression networks. Statistical comparison of the degree of genes between SD patients and healthy controls yielded a total of 647 Differentially Interacting Genes (DIGs) at a false discovery rate of 5% (*p*_*adj*_ < 0.05). The full list of DIGs, including their degree differences and adjusted p-values, is available as Supplementary Table S1.

The global distribution of these topological changes, as represented in the volcano plot (Figure 1, Panel A), indicates a predominant gain of connectivity within the SD cohort. Several well-characterized interferon-stimulated genes (ISGs) emerged as the most significant topological hubs. Specifically, *USP18* (Δ*Degree* = 14.88), *IFI44L* (Δ*Degree* = 14.39), and *SIGLEC1* (Δ*Degree* = 24.14) displayed the most substantial increases in interaction degree.

**Figure 1:**
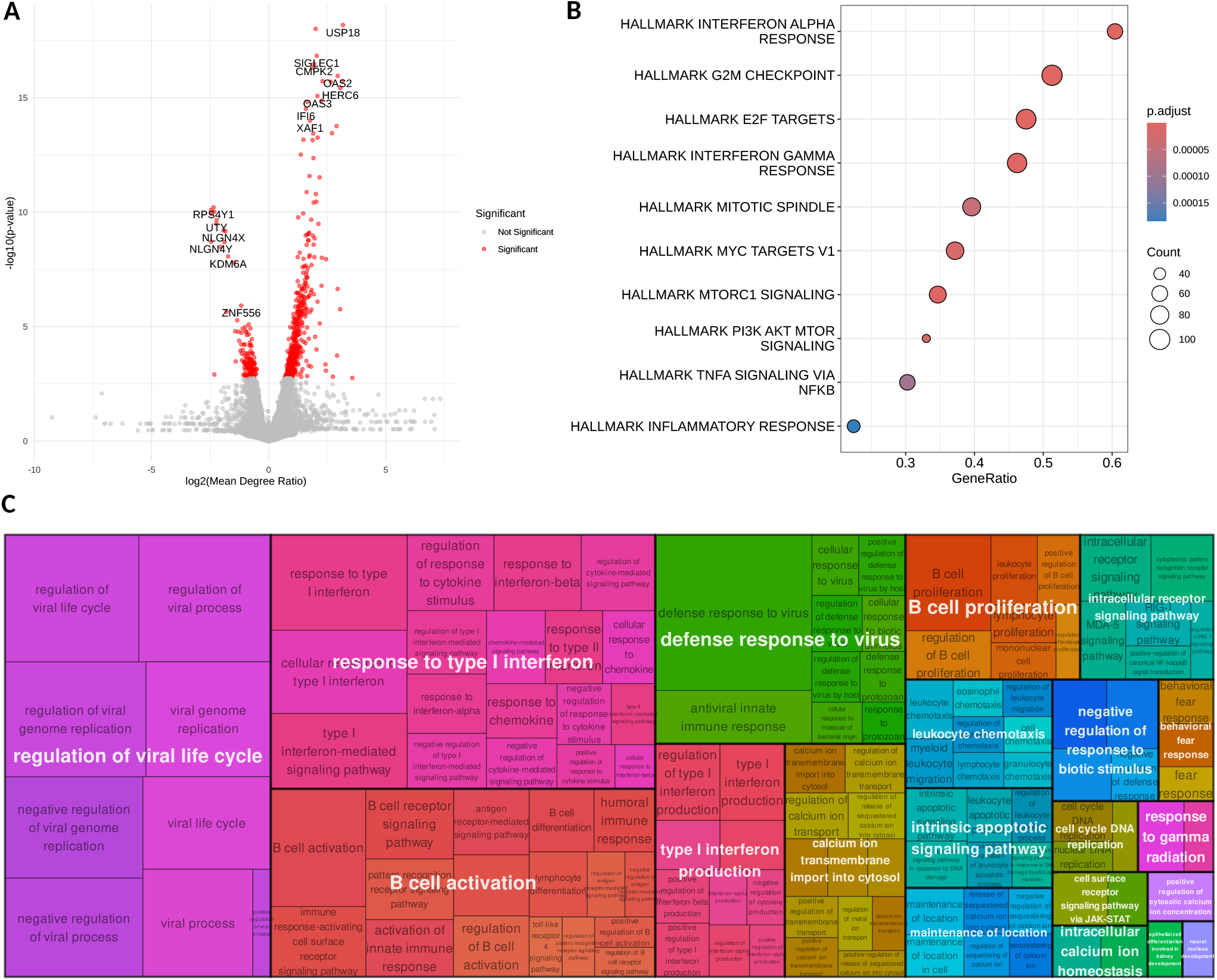
Global landscape of differential gene interactivity in SD. **(A)** Volcano plot illustrating the distribution of 647 Differentially Interacting Genes (DIGs) identified via LIONESS between SD patients and healthy controls. The y-axis represents − log_10_(p-value) and the x-axis shows the log_2_ ratio of mean connectivity (degree). Red points indicate genes meeting the significance threshold (*p*_*adj*_ < 0.05); top-ranked topological hubs are highlighted. **(B)** Bar chart of the top 10 most significant GO biological processes enriched for the DIGs with an increase in degree in the comparison.**(C)** Hierarchical treemap of GO biological processes enriched among the 647 DIGs. The size of each rectangle corresponds to the − log_10_(p-value) of the enrichment.

To determine the biological processes governed by this rewired network, we performed both Gene Set Enrichment Analysis using Hallmark processes (Figure 1, Panel B) as well as overrepresentation analysis using Gene Ontology (GO) on the 647 identified DIGs. The functional landscape of the overrepresentation analysis is summarized in the GO treemap (Figure 1, Panel C). The most robustly enriched biological processes were centered on response to type I and type II interferons, including terms such as response to interferon-alpha and interferon-gamma-mediated signaling. Additionally, significant enrichment was observed in antiviral defense mechanisms and nucleic acid sensing, as well as antigen processing and presentation.

To contextualize the higher-order structural organization of the 647 identified DIGs, we constructed a physical protein-protein interaction (PPI) network using experimentally validated links from the STRING database. Community detection using the Louvain algorithm resolved the network into distinct topological modules. By collapsing interconnected sub-communities sharing redundant Gene Ontology (GO) terms, we identified four major functional macro-modules: *defense response to virus, B cell activation, DNA replication*, and *positive regulation of protein catabolic process* (Figure 2). The major hubs within each cluster were ranked by their sample-specific degree, mapping absolute connectivity changes (|ΔDegree|) directly to node size.

**Figure 2:**
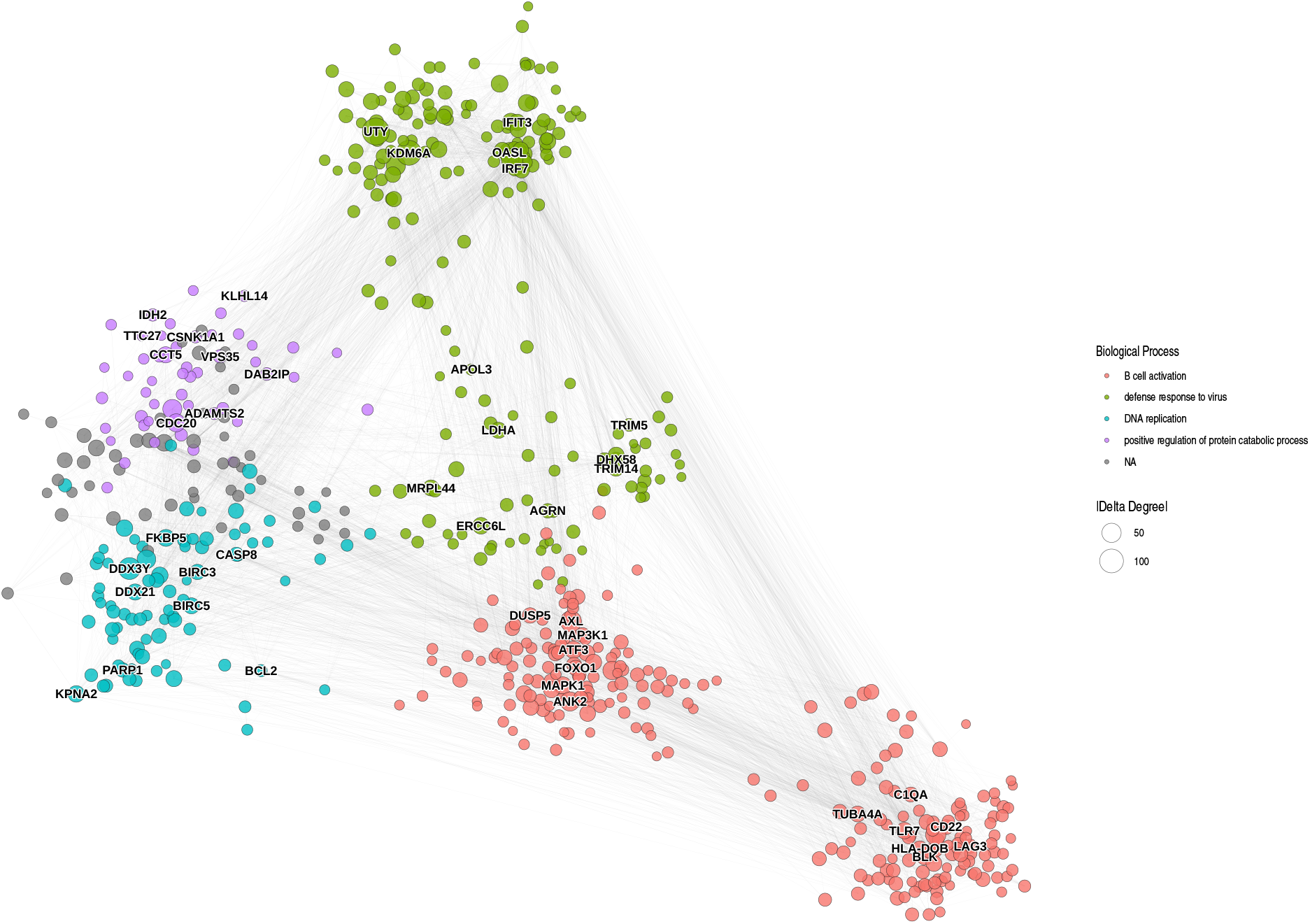
Protein-protein interaction network of differentially interacting genes in SD. The network illustrates experimentally validated physical interactions among the significant Differentially Interacting Genes (DIGs) identified between SD patients and healthy controls. Nodes represent individual DIGs, with node size proportional to the absolute change in connectivity (|ΔDegree |). Nodes are clustered and colored according to their primary Gene Ontology (GO) biological process, identified through community clustering and subsequent functional enrichment. The top 10 topological hubs within each major functional cluster (ranked by degree) are annotated with their gene symbols. Edges denote protein-protein interactions, displayed using a layout that emphasizes the network’s structural backbone.

The largest macro-module identified within the rewired network belongs to the *defense response to virus* process (olive cluster, Figure 2). This cluster is dominated by a dense core of classic interferon-stimulated genes (ISGs). Notably, *ISG15* emerged as the single most rewired topological hub in the entire dataset (|ΔDegree| = 75.10). It is accompanied by heavily characterized clinical biomarkers used for patient stratification, including *OASL* (|ΔDegree| = 61.84), *IFIT1* (| ΔDegree| = 31.69), *IFIT3* (|ΔDegree| = 28.14), and *MX1* (|ΔDegree| = 24.90). Cytosolic nucleic acid sensors such as *RIGI* (*DDX58*, |ΔDegree| = 27.82) and *IFIH1* (|ΔDegree| = 24 .51) anchor the interior of this module.

The second prominent macro-module encompasses processes driving *B cell activation* (coral cluster, Figure 2). *AXL*, a member of the TAM receptor tyrosine kinase family, exhibits a massive gain of baseline connectivity (|ΔDegree| = 68.54), placing it as the major coordinator of this module. Crucially, the myeloid-specific marker *SIGLEC1* (*CD169*, |ΔDegree| = (24.15) is topologically embedded within this adaptive immune cluster.

The third macro-module centers on *DNA replication* and cell cycle proliferation (green cluster, Figure 2). Highly connected hubs inside this cluster include the mitotic regulators *TPX2* (|ΔDegree| = 11.92) and *BIRC5* (*Survivin*, |ΔDegree| = (25.11), alongside the proliferation anchor *MKI67* (|ΔDegree|= 12 61).

While our DIG network successfully captured canonical, heavily documented genes in SD pathogenesis, it simultaneously brought to light several highly rewired hubs that represent major gaps in current disease literature. Within the antiviral defense module, the ISGylation mediator *UBE2L6* (|ΔDegree| = 21.97) and the RNA-binding protein *IFIT5* (|ΔDegree| = 30.95) emerged as core hubs despite a lack of functional validation in Sjögren’s cohorts. Similarly, the early B cell receptor development marker *VPREB1* (|ΔDegree| = 6.63) and the signaling mediator *PRKCE* (|ΔDegree| = 14.97) were heavily integrated into the B cell activation module.

To further establish the biological relevance of the identified DIGs, we integrated individual gene connectivity metrics with biological process activity scores derived from ssGSEA. This analysis aimed to determine if the interaction degree of a gene scales proportionally with the intensity of the biological process it belongs to.

Spearman correlation analysis demonstrated a high degree of synchronization between the connectivity of primary hubs and their respective pathways. For instance, the interaction degrees of the network hubs *OASL, UBE2L6, ISG15* and *LGALS3BP* showed strong positive correlations (ρ > 0.4, *p* < 0.001) with the Defense Response to Virus scores across the cohort (Figure 3).

**Figure 3:**
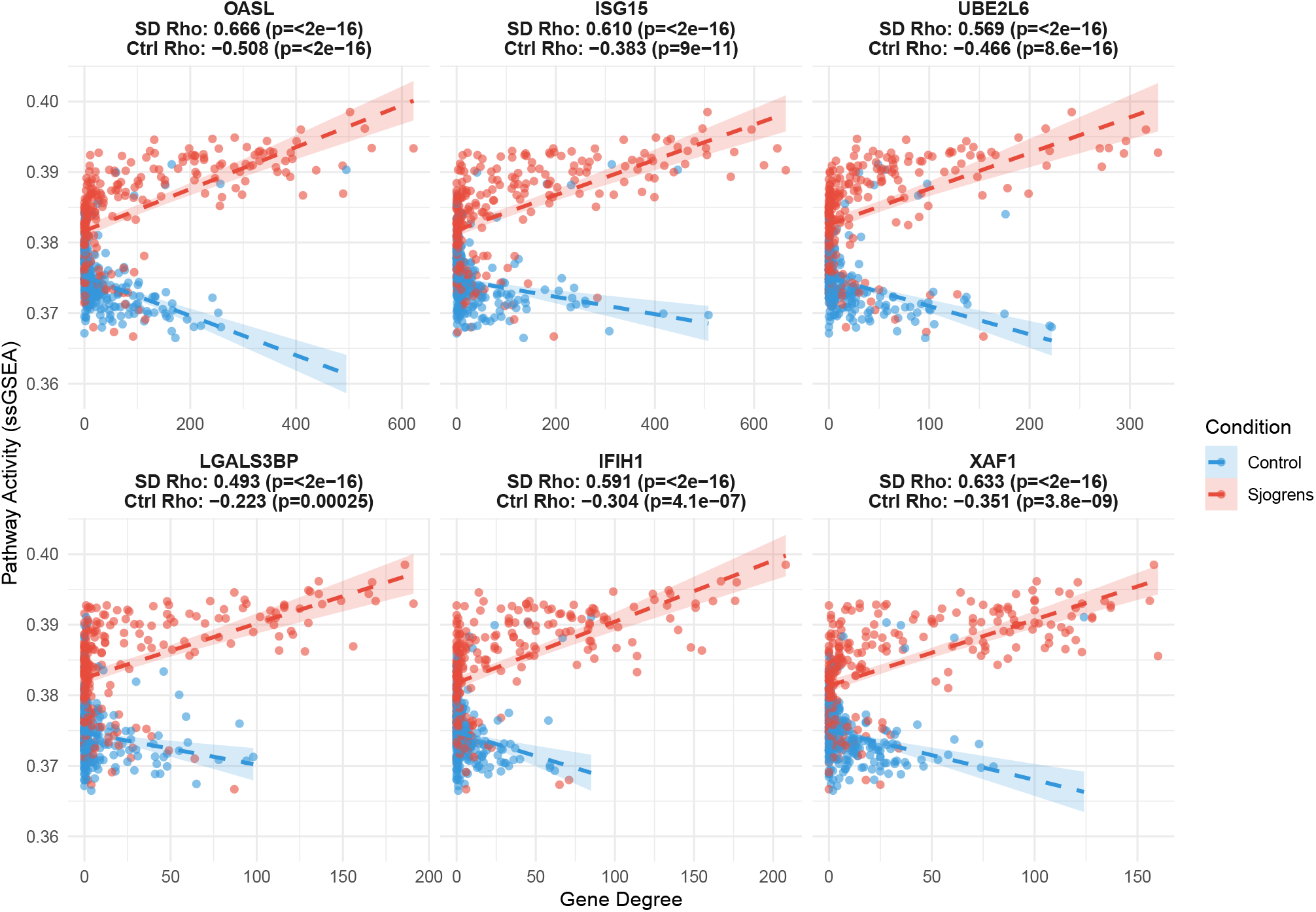
Correlation between sample-specific gene degree and GOBP_DEFENSE_RESPONSE_TO_VIRUS pathway activity in Sjögren’s disease and healthy controls. Scatter plots illustrate the coupling between individual gene interaction connectivity (sample-specific degree, x-axis) and overall pathway activity derived via ssGSEA (y-axis) for the *GOBP_DEFENSE_RESPONSE_TO_VIRUS* signature. Each panel displays a highly rewired topological hub: **(A)** *OASL*, **(B)** *ISG15*, **(C)** *UBE2L6*, **(D)** *LGALS3BP*, **(E)** *IFIH1*, and **(F)** *XAF1*. Individual subjects are stratified by clinical status into healthy controls (blue) and Sjögren’s disease patients (red). To accurately visualize the reported Spearman rank correlations (ρ), dashed trendlines were fitted using a Robust Linear Model (RLM), with shaded areas representing 95% confidence intervals. Spearman rank correlations are reported separately for each group; note that the sign of the correlation reverses between patients (ρ = +0.49 to +0.67) and controls (ρ = −0.22 to −0.51) for every gene shown.

To identify the most critical structural vulnerabilities within this dysregulated network, we performed an in-silico knockout analysis on the patient-specific weighted network, measuring the fractional drop in Global Efficiency following removal of each druggable DIG (Table 2). *EPHB2* ranked highest and also showed the largest degree-corrected residual, indicating a structural contribution well in excess of its connectivity, followed by *BLK* and *SERPING1*. Several candidates of low degree ranked highly once connectivity was accounted for, including *CYSLTR1, BLVRA* and *NT5E. MAPK1* ranked fourth, but its centrality reflects its role as a general signalling hub and the approved compounds annotated against it are non-selective. The complete network vulnerability results for all druggable DIGs are available as Supplementary Table S2.

**Table (2.** Top 15 druggable Differentially Interacting Genes ranked by the fractional drop in Global Efficiency (GE) following in-silico deletion from the patient-specific network. Vulnerability is reported for the networks weighted by the median LIONESS edge weight across SD patients. The residual column gives the deviation from a linear regression of vulnerability on node degree, isolating the contribution not explained by connectivity alone. Representative drugs are approved agents annotated against each target in DrugBank, except Conestat alfa, a recombinant C1-inhibitor annotated against *SERPING1* in ChEMBL, and Epratuzumab, an investigational anti-CD22 antibody previously evaluated in SD. All approved DrugBank drugs with an annotated action on each target are listed in Supplementary Table S2.

| Target | Degree | GE Vuln. | Degree-corr. residual | Representative Drug |
| --- | --- | --- | --- | --- |
| *EPHB2* | 96 | 0.00625 | +0.00532 | Fostamatinib |
| *BLK* | 174 | 0.00444 | +0.00209 | Zanubrutinib |
| *SERPING1* | 34 | 0.00419 | +0.00439 | Conestat alfa |
| *MAPK1* | 186 | 0.00380 | +0.00123 | Minocycline |
| *TLR7* | 158 | 0.00233 | +0.00027 | Hydroxychloroquine |
| *CYSLTR1* | 12 | 0.00144 | +0.00204 | Montelukast |
| *TUBA4A* | 118 | 0.00138 | +0.00005 | Vincristine |
| *CD22* | 110 | 0.00132 | +0.00013 | Epratuzumab |
| *AXL* | 140 | 0.00132 | -0.00042 | Gilteritinib |
| *NT5E* | 40 | 0.00126 | +0.00135 | Theophylline |
| *PARP1* | 80 | 0.00102 | +0.00038 | Cladribine |
| *BLVRA* | 8 | 0.00081 | +0.00148 | Ursodeoxycholic acid |
| *CAMK2G* | 68 | 0.00079 | +0.00037 | Bosutinib |
| *MAP3K1* | 114 | 0.00073 | -0.00053 | Fostamatinib |
| *CTSL* | 44 | 0.00067 | +0.00069 | Fostamatinib |

This vulnerability assessment highlighted a highly druggable signaling landscape dominated by kinases. *BLK, EPHB2, MAPK1*, and *AXL* dominated the top rankings. Knocking out these targets forced communication pathways between the remaining nodes to reroute through significantly longer topological paths. The endosomal receptor *TLR7* and the lysosomal protease *CTSL* also emerged as highly efficient routing bottlenecks.

## 4. Discussion

The molecular transition from health to Sjögren’s Disease (SD) is characterized by a fundamental shift in how gene products coordinate their activity. By employing sample-specific network modeling, our analysis captures the dynamic rewiring of the interactome. Previous studies have demonstrated that network propagation and topological modeling can act as universal amplifiers of genetic associations, uncovering disease modules that are otherwise obscured in standard expression analyses Cowen et al. (2017); Scelsi et al. (2021). Our findings suggest that the inflammatory state in SD promotes a large-scale topological transition, where specific genes move from the network periphery to become central regulatory anchors, effectively stabilizing the chronic autoimmune response.

The massive topological recruitment of these genes demonstrates that the well-characterized “interferon signature” of SD is underpinned by a profound structural reorganization. While the systemic upregulation of type I and type II interferons (IFN) is a well-established driver of SD pathogenesis Bodewes et al. (2018); Gupta et al. (2024), our network reveals that classical interferon-stimulated genes (ISGs) such as *ISG15, USP18*, and *IFI44L* undergo a robust gain in connectivity. The high degree of synchronization between the connectivity of these primary hubs and their respective pathway activity scores indicates that these DIGs act as functional anchors. As the pathological activity of the interferon pathway increases, these genes structurally recruit additional interaction partners to reinforce the transcriptional program.

In this context, the topological recruitment of genes into these modules suggests a coordinated mechanism by which the innate interferon signal is structurally linked to the adaptive immune dysfunction observed in SD patients. Our network delineates an active innate-to-adaptive bridge in the tissue environment: rewired antigen-presenting cells expressing the myeloid marker *SIGLEC1*, a known biomarker of SD disease activity and extraglandular manifestation Fox et al. (2021), and signaling through the TAM receptor *AXL* are physically and topologically positioned to sustain the survival and clonal expansion of infiltrating B cell subsets. While deficiencies or alterations in TAM receptor signaling (Tyro3, Axl, and Mer) have been associated with impaired efferocytosis and hyperproliferation of lymphocytes in autoimmune conditions Choi et al. (2025), the massive gain of baseline connectivity for *AXL* (|ΔDegree| = 68.54) places it as a major coordinator of this adaptive cluster.

This architecture supports the emergence of a prosurvival and anti-apoptotic module in B cells. B cell hyperactivity and the development of pathogenic autoreactive clones are central to SD Verstappen et al. (2021); Du et al. (2021). Our data indicate that this hyperactivity is structurally driven by signaling axes involving *CXCR4, PRKCE*, and *VPREB1. CXCR4* is known to play a pivotal role in leukocyte trafficking and B-cell survival Schweighoffer and Tybulewicz (2018); Giorgiutti et al. (2024), and its high connectivity within this module likely underpins the aberrant costimulation and prolonged survival of B cells in the glandular microenvironment.

Furthermore, the data indicate that while chronic tissue damage restricts normal epithelial regeneration, infiltrating autoreactive immune clones rely on specific, rewired cell cycle hubs to evade apoptosis and maintain a hyperproliferative state. Most notably, the cell cycle replication and apoptosisevasion anchors *BIRC5* (Survivin), *FEN1* (|ΔDegree| = 17.94), and *TPX2* constitute novel discoveries in this context. The high network centrality of these previously uncharacterized genes suggests they represent critical, unrecognized structural nodes driving disease pathogenesis.

By interrogating the structural integrity of this dysregulated network, we identified vulnerabilities spanning a few targets of interest.

Among them is *CD22*, which has the most supporting literature at the moment. Epratuzumab, a humanised anti-CD22 antibody, was evaluated in an open-label study in primary SD, where a composite response was achieved by a majority of patients Steinfeld et al. (2006). In a subsequent post hoc analysis of the EMBODY trials in systemic lupus erythematosus, the subgroup of anti-SSA-positive patients with associated Sjögren’s syndrome showed greater clinical response and deeper B-cell depletion than lupus patients without it Gottenberg et al. (2018). *CD19*, likewise recovered by the analysis, is the subject of an ongoing trial in SD, and anti-CD19 therapy has produced serological remission in a patient with concurrent SD and lymphoma Sheng et al. (2023) as well as efficacy in IgG4-related disease Stone et al. (2025).

*EPHB2* represents the clearest novel prediction. It ranked first in the vulnerability analysis and showed the largest degree-corrected residual of any candidate, yet has no published description in SD. EphB2 knockdown has been shown to reduce proliferation, TNF-α production and immunoglobulin secretion in human naive B cells Yu et al. (2014), and anti-EphB2 autoantibodies are detectable in lupus and scleroderma Azzouz et al. (2016), although in the latter case as an autoantigen rather than a signalling driver.

Two further candidates warrant mention. *CYSLTR1* was independently identified as an overexpressed diagnostic marker in SD by an expression-based pipeline, where it performed comparably to *TNFSF13B* Zeng et al. (2022); the receptor is expressed in the human ocular surface Brunner et al. (2023), and montelukast limits acinar atrophy and fibrosis in salivary and lacrimal glands in rodent injury models Koca et al. (2013a,b) and modulates Th17 differentiation in experimental autoimmune encephalomyelitis Han et al. (2021). *SERPING1* encodes the principal regulator of the classical complement pathway and is interferon-inducible Hall et al. (2012); complement consumption is an established prognostic marker in SD, with C4 hypocomplementaemia independently predicting lymphoma Fragkioudaki et al. (2016), and C1-inhibitor deficiency co-occurs with SD Triggianese et al. (2024). Direct evidence implicating *SERPING1* itself, however, is limited to a proteomic study in an experimental model Li et al. (2021).

While this in-silico topological framework offers insights into SD pathogenesis, several limitations must be acknowledged. First, the reconstructed networks rely on bulk RNA sequencing. Although this captures the systemic immune signature, it lacks the fine spatial and cellular resolution achieved in recent single-cell transcriptomic atlases of SD, which are necessary to pinpoint whether specific interactions occur exclusively within distinct cellular sub-populations (e.g., specific CD8+ T cells or ductal epithelial cells) Xiang et al. (2023); Pranzatelli et al. (2024). Second, the underlying dataset consisted of single-end short reads. This sequencing architecture inherently restricts the ability to capture complex alternative splicing events or isoform-specific network rewiring that could further define SD pathogenesis. Third, the restriction of edges to those experimentally validated in the STRING database ensures biological plausibility but limits the discovery of entirely novel, context-specific interactions. Finally, while the topological repositioning of FDA-approved drugs provides strong computational validation for novel targets, these candidates strictly require subsequent *in vitro* and *in vivo* functional validation to confirm their therapeutic efficacy and safety.

In conclusion, our single-sample network analysis successfully reconstructed key signaling pathways dysregulated in SD, demonstrating that disease pathogenesis is driven by deep structural rewiring of the interactome. By integrating network topology with functional pathway activity, we identified novel diagnostic anchors and characterized a structurally distinct innate-to-adaptive immune bridge driving B-cell hyperactivity. Ultimately, this approach uncovers previously obscured, druggable topological vulnerabilities, providing a computationally robust framework for future therapeutic development and personalized drug repositioning in Sjögren’s disease.

## Availability of data and materials

The code used in this study is available at https://github.com/mcblab/sd_nets. PRECISESADS data is hosted by ELIXIR Luxembourg. The dataset is available for research use under a controlled access model. To request access please contact the data stewardship team of ELIXIR Luxembourg via.

## Funding

This study was financed by the Brazilian Agency Coordination for the Improvement of Higher Education Personnel (CAPES – Portuguese: Coordenação de Aperfeiçoamento de Pessoal de Nível Superior - Brasil (CAPES)- (project numbers 88881.692896/2022-01 and 88887.002655/2024-00), National Council of Technological and Scientific Development (CNPq – Portuguese: Conselho Nacional de Desenvolvimento Científico e Tecnológico (CNPq) (project number 312305/2021-4 and 305019/2025-2), PROPESQ-UFRN.

## Acknowledgements

The authors would like to thank NPAD / UFRN for computational resources.

## CRediT authorship contribution statement

**João V. F. Cavalcante:** Conceptualization, Formal analysis, Investigation, Methodology, Visualization, Writing – original draft, Writing – review and editing. **Rodrigo J. S. Dalmolin:** Supervision, Writing – review and editing. **Diego Marques-Coelho:** Conceptualization, Data curation, Investigation, Writing – review and editing.

## Notes

### Competing Interest Statement

The authors have declared no competing interest.

## References

Azzouz, D.F., et al., 2016. Anti-ephrin type-B receptor 2 (EphB2) and anti-three prime histone mRNA EXonuclease 1 (THEX1) autoantibodies in scleroderma and lupus. PLoS ONE 11, e0160283. doi:10.1371/journal.pone.0160283. pMID: 27617966. EphB2 as autoantigen, not as a signalling driver.

Barturen, G., Babaei, S., Català-Moll, F., Martínez-Bueno, M., Makowska, Z., Martorell-Marugán, J., Carmona-Sáez, P., Toro-Domínguez, D., Carnero-Montoro, E., Teruel, M., Kerick, M., Acosta-Herrera, M., Lann, L.L., Jamin, C., Rodríguez-Ubreva, J., García-Gómez, A., Kageyama, J., Buttgereit, A., Hayat, S., Mueller, J., Lesche, R., Hernandez-Fuentes, M., Juarez, M., Rowley, T., White, I., Marañón, C., Anjos, T.G., Varela, N., Aguilar-Quesada, R., Garrancho, F.J., López-Berrio, A., Maresca, M.R., Navarro-Linares, H., Almeida, I., Azevedo, N., Brandão, M., Campar, A., Faria, R., Farinha, F., Marinho, A., Neves, E., Tavares, A., Vasconcelos, C., Trombetta, E., Montanelli, G., Vigone, B., Alvarez-Errico, D., Li, T., Thiagaran, D., Alonso, R.B., Martínez, A.C., Genre, F., Mejías, R.L., Gonzalez-Gay, M.A., Remuzgo, S., Garcia, B.U., Cervera, R., Espinosa, G., Rodríguez-Pintó, I., Langhe, E.D., Cremer, J., Lories, R., Belz, D., Hunzelmann, N., Baerlecken, N., Kniesch, K., Witte, T., Lehner, M., Stummvoll, G., Zauner, M., Aguirre-Zamorano, M.A., Barbarroja, N., Castro-Villegas, M.C., Collantes-Estevez, E., de Ramon, E., Quintero, I.D., Escudero-Contreras, A., Roldán, M.C.F., Gómez, Y.J., Moleón, I.J., Lopez-Pedrera, R., Ortega-Castro, R., Ortego, N., Raya, E., Artusi, C., Gerosa, M., Meroni, P.L., Schioppo, T., Groof, A.D., Ducreux, J., Lauwerys, B., Maudoux, A.L., Cornec, D., Devauchelle-Pensec, V., Jousse-Joulin, S., Jouve, P.E., Rouvière, B., Saraux, A., Simon, Q., Alvarez, M., Chizzolini, C., Dufour, A., Wynar, D., Balog, A., Bocskai, M., Deák, M., Dulic, S., Kádár, G., Kovács, L., Cheng, Q., Gerl, V., Hiepe, F., Khodadadi, L., Thiel, S., de Rinaldis, E., Rao, S., Benschop, R.J., Chamberlain, C., Dow, E.R., Ioannou, Y., Laigle, L., Marovac, J., Wojcik, J., Renaudineau, Y., Borghi, M.O., Frostegård, J., Martín, J., Beretta, L., Ballestar, E., McDonald, F., Pers, J.O., Alarcón-Riquelme, M.E., 2021. Integrative Analysis Reveals a Molecular Stratification of Systemic Autoimmune Diseases. Arthritis & Rheumatology 73, 1073–1085. doi:10.1002/art.41610.

Bodewes, I.L.A., Huijser, E., van Helden-Meeuwsen, C.G., Tas, L., Huizinga, R., Dalm, V.A.S.H., van Hagen, P.M., Groot, N., Kamphuis, S., van Daele, P.L.A., Versnel, M.A., 2018. TBK1: A key regulator and potential treatment target for interferon positive Sjögren’s syndrome, systemic lupus erythematosus and systemic sclerosis. Journal of Autoimmunity 91, 97–102. doi:10.1016/j.jaut.2018.02.001.

Bowman, S.J., Fox, R., Dörner, T., et al., 2022. Safety and efficacy of sub-cutaneous ianalumab (VAY736) in patients with primary Sjögren’s syndrome: A randomised, double-blind, placebo-controlled, phase (2b dosefinding trial. The Lancet 399, 161–171. doi:10.1016/S0140-673621)02251-0. pMID: 34861168. Phase 3 NEPTUNUS-1/-2 both met their primary ESSDAI endpoints; FDA Breakthrough designation January 2026.

Brunner, S.M., Schrödl, F., Preishuber-Pflügl, J., et al., 2023. Distribution of the cysteinyl leukotriene system components in the human, rat and mouse eye. Experimental Eye Research 232, 109517. doi:10.1016/j.exer.2023.109517. pMID: 37211287. Confirms CysLTR1 is present in human ocular surface tissue.

Carlson, M., . Org.Hs.eg.db. doi:10.18129/B9.bioc.org.Hs.eg.db.

Chen, H.H., Hsueh, C.W., Lee, C.H., Hao, T.Y., Tu, T.Y., Chang, L.Y., Lee, J.C., Lin, C.Y., 2023. SWEET: A single-sample network inference method for deciphering individual features in disease. Briefings in Bioinformatics 24, bbad032. doi:10.1093/bib/bbad032.

Choi, S.E., Kang, J.H., Park, D.J., Kim, N.I., Lee, J.S., Yoon, K.C., Lee, S.S., 2025. Clinical significance of TAM receptor in the minor salivary glands of patients with Sjögren’s disease. Scientific Reports 15, 23065. doi:10.1038/s41598-025-08086-z.

Cowen, L., Ideker, T., Raphael, B.J., Sharan, R., 2017. Network propagation: A universal amplifier of genetic associations. Nature Reviews Genetics 18, 551–562. doi:10.1038/nrg.2017.38.

Deschildre, J., Vandemoortele, B., Loers, J.U., De Preter, K., Vermeirssen, V., 2024. Evaluation of single-sample network inference methods for precision oncology. npj Systems Biology and Applications 10, 1–16. doi:10.1038/s41540-024-00340-w.

Du, W., Han, M., Zhu, X., Xiao, F., Huang, E., Che, N., Tang, X., Zou, H., Jiang, Q., Lu, L., 2021. The Multiple Roles of B Cells in the Pathogenesis of Sjögren’s Syndrome. Frontiers in Immunology 12. doi:10.3389/fimmu.2021.684999.

Fan, P., Kofler, J., Ding, Y., Marks, M., Sweet, R.A., Wang, L., 2022. Efficacy difference of antipsychotics in alzheimer’s disease and schizophrenia: explained with network efficiency and pathway analysis methods. Briefings in Bioinformatics 23, bbac394. URL: https://doi.org/10.1093/bib/bbac394, doi:10.1093/bib/bbac394.

Fisher, B.A., et al., 2024. Effect of iscalimab on disease activity in patients with Sjögren’s disease (TWINSS): A randomised, doubleblind, placebo-controlled, phase 2b trial. The Lancet 404, 540–553. doi:10.1016/S0140-6736(24)01211-X. pMID: 39096929. Cohort 1 met the ESSDAI dose-response endpoint; cohort 2 (low ESSDAI / high symptom burden) was negative on ESSPRI.

Fox, R.I., Fox, C.M., Gottenberg, J.E., Dörner, T., 2021. Treatment of Sjögren’s syndrome: Current therapy and future directions. Rheumatology 60, 2066–2074. doi:10.1093/rheumatology/kez142.

Fragkioudaki, S., Mavragani, C.P., Moutsopoulos, H.M., 2016. Predicting the risk for lymphoma development in Sjögren syndrome: An easy tool for clinical use. Medicine (Baltimore) 95, e3766. doi:10.1097/MD.0000000000003766. pMID: 27336863. C4 hypocomplementemia is an independent predictor of non-Hodgkin lymphoma (381 cases vs 92 controls).

Giorgiutti, S., Rottura, J., Korganow, A.S., Gies, V., 2024. CXCR4: From B-cell development to B cell–mediated diseases. Life Science Alliance 7, e202302465. doi:10.26508/lsa.202302465.

Gottenberg, J.E., Dörner, T., Bootsma, H., Devauchelle-Pensec, V., Bowman, S.J., Mariette, X., 2018. Efficacy of epratuzumab, an anti-CD22 monoclonal IgG antibody, in systemic lupus erythematosus patients with associated Sjögren’s syndrome: Post hoc analyses from the EMBODY trials. Arthritis & Rheumatology 70, 763–773. doi:10.1002/art.40425. pMID: 29381843. The 113 anti-SSA+ patients with associated Sjögren’s showed higher BICLA response and deeper B-cell depletion than SLE patients without SS.

Gupta, S., Yamada, E., Nakamura, H., Perez, P., Pranzatelli, T.J., Dominick, K., Jang, S.I., Abed, M., Martin, D., Burbelo, P., Zheng, C., French, B., Alevizos, I., Khavandgar, Z., Beach, M., Pelayo, E., Walitt, B., Hasni, S., Kaplan, M.J., Tandon, M., Magone, M.T., Kleiner, D.E., Chiorini, J.A., Baer, A., Warner, B.M., 2024. Inhibition of JAK-STAT pathway corrects salivary gland inflammation and interferon driven immune activation in Sjögren’s disease. Annals of the Rheumatic Diseases 83, 1034–1047. doi:10.1136/ard-2023-224842.

Hall, J.C., Casciola-Rosen, L., Kapsogeorgou, E.K., Tzioufas, A.G., Baer, A.N., Rosen, A., 2012. Precise probes of type II interferon activity define the origin of interferon signatures in target tissues in rheumatic diseases. Proceedings of the National Academy of Sciences 109, 17609–17614. doi:10.1073/pnas.1209724109. pMID: 23045702. SERPING1 is part of the interferon probe gene set.

Han, B., Zhang, Y.Y., Ye, Z.Q., et al., 2021. Montelukast alleviates inflammation in experimental autoimmune encephalomyelitis by altering Th17 differentiation in a mouse model. Immunology 163, 185–200. doi:10.1111/imm.13312. pMID: 33480040.

Hänzelmann, S., Castelo, R., Guinney, J., 2013. GSVA: Gene set variation analysis for microarray and RNA-Seq data. BMC Bioinformatics 14, 7. doi:10.1186/1471-2105-14-7.

Koca, G., Gültekin, S.S., Han, U., Kuru, S., Demirel, K., Korkmaz, M., 2013a. The efficacy of montelukast as a protective agent against 131I-induced salivary gland damage in rats: Scintigraphic and histopathological findings. Nuclear Medicine Communications 34, 507–516. doi:10.1097/MNM.0b013e32835ffecd. pMID: 23478587.

Koca, G., Yalniz-Akkaya, Z., Gültekin, S.S., et al., 2013b. Radioprotective effect of montelukast sodium in rat lacrimal glands after radioiodine treatment. Revista Española de Medicina Nuclear e Imagen Molecular 32, 240–245. PMID: 23499122. Reduced acinar atrophy and fibrosis in intraorbital, extraorbital and Harderian glands.

Koutsandreas, T., Tsafou, K., Horn, H., Barrett, I., Petsalaki, E., 2025. Network-Based Approaches for Drug Target Identification. Annual Review of Biomedical Data Science 8. doi:10.1146/annurev-biodatasci-101424-120950.

Kuijjer, M.L., Hsieh, P.H., Quackenbush, J., Glass, K., 2019a. lionessR: Single sample network inference in R. BMC Cancer 19, 1003. doi:10.1186/s12885-019-6235-7.

Kuijjer, M.L., Tung, M.G., Yuan, G., Quackenbush, J., Glass, K., 2019b. Estimating Sample-Specific Regulatory Networks. iScience 14, 226–240. doi:10.1016/j.isci.2019.03.021.

Laigle, L., Beretta, L., Wojick, J., Marovac, J., Pers, J.O., Lauwerys, B., Frostegard, J., Donald, F.M., Juarez, M., Benschop, R., Dow, E., Rao, S., Chamberlain, C., Martin, J., Alarcon-Riquelme, M.E., Consortium, o.b.o.t.C., 2018. AB1372 Towards reforming the taxonomy of human disease: The precisesads cross sectional study. Annals of the Rheumatic Diseases 77, 1771–1772. doi:10.1136/annrheumdis-2018-eular.4625.

Li, M., Qi, Y., Wang, G., et al., 2021. Proteomic profiling of saliva reveals association of complement system with primary Sjögren’s syndrome. Immunity, Inflammation and Disease 9. doi:10.1002/iid3.529. pMID: 34516718. SERPING1 downregulated in saliva and a core PPI node. IMPORTANT CAVEAT: experimental SS mouse model, n=3 vs 3, not human patients.

Liberzon, A., Birger, C., Thorvaldsdóttir, H., Ghandi, M., Mesirov, J.P., Tamayo, P., 2015. The Molecular Signatures Database (MSigDB) hallmark gene set collection. Cell systems 1, 417–425. doi:10.1016/j.cels.2015.12.004.

López-Sánchez, P., Ávila-Moreno, F., Hernández-Lemus, E., Kuijjer, M.L., Espinal-Enríquez, J., 2025. Patient-specific gene co-expression networks reveal novel subtypes and predictive biomarkers in lung adenocarcinoma. npj Systems Biology and Applications 11, 44. doi:10.1038/s41540-025-00522-0.

Mavragani, C.P., Moutsopoulos, H.M., 2020. Sjögren’s syndrome: Old and new therapeutic targets. Journal of Autoimmunity 110, 102364. doi:10.1016/j.jaut.2019.102364.

Noaiseh, G., et al., 2025. Nipocalimab in primary Sjögren’s disease (DAHLIAS): A multicentre, double-blind, randomised, placebocontrolled, phase (2 study. The Lancet 406, 2435–2448. doi:10.1016/S0140-6736(25)01430-8. pMID: 41284548.

Pranzatelli, T.J., Perez, P., Ku, A., Matuck, B.F., Huynh, K., Sakai, S., Abed, M., Jang, S.I., Yamada, E., Dominick, K., Ahmed, Z., Oliver, A., Wasikowski, R., Easter, Q.T., Baer, A.N., Pelayo, E., Khavandgar, Z., Gupta, S., Kleiner, D.E., Magone, M.T., Lessard, C., Farris, A.D., Burbelo, P.D., Martin, D., Morell, R., Zheng, C., Rachmaninoff, N., Maldonado-Ortiz, J., Qu, X., Aure, M.H., Dezfulian, M.H., Lake, R., Teichmann, S., Barber, D.L., Tsoi, L.C., Sowalsky, A.G., Tyc, K.M., Liu, J., Gudjonsson, J.E., Byrd, K.M., Johnson, P.L., Chiorini, J.A., Warner, B.M., 2024. GZMK+CD8+ T cells Target A Specific Acinar Cell Type in Sjögren’s Disease. doi:10.21203/rs.3.rs-3601404/v2.

Ritchie, M.E., Phipson, B., Wu, D., Hu, Y., Law, C.W., Shi, W., Smyth, G.K., 2015. Limma powers differential expression analyses for RNA-sequencing and microarray studies. Nucleic Acids Research 43, e47. doi:10.1093/nar/gkv007.

Scelsi, M.A., Napolioni, V., Greicius, M.D., Altmann, A., Project (ADSP), f.t.A.D.N.I.A.a.t.A.D.S., 2021. Network propagation of rare variants in Alzheimer’s disease reveals tissue-specific hub genes and communities. PLOS Computational Biology 17, e1008517. doi:10.1371/journal.pcbi.1008517.

Schweighoffer, E., Tybulewicz, V.L., 2018. Signalling for B cell survival. Current Opinion in Cell Biology 51, 8–14. doi:10.1016/j.ceb.2017.10.002.

Sheng, L., Zhang, Y., Song, Q., et al., 2023. Concurrent remission of lymphoma and Sjögren’s disease following anti-CD19 chimeric antigen receptor-T cell therapy for diffuse large B-cell lymphoma: A case report. Frontiers in Immunology 14, 1298815. doi:10.3389/fimmu.2023.1298815. anti-Ro52 and ANA became negative at day 90; ESSDAI 5 to 2. CAR-T was given for lymphoma, not for SjD.

St Clair, E.W., et al., 2024. Efficacy and safety of dazodalibep in Sjögren’s disease: A randomised, double-blind, placebo-controlled, phase 2 trial. Nature Medicine 30, 1583–1592. doi:10.1038/s41591-024-03009-3. pMID: 38839899.

Steinfeld, S.D., Tant, L., Burmester, G.R., et al., 2006. Epratuzumab (humanised anti-CD22 antibody) in primary Sjögren’s syndrome: An open-label phase I/II study. Arthritis Research & Therapy 8, R129. doi:10.1186/ar2018. pMID: 16859536. n=16, 360 mg/m2 x4. Composite response 53% at week 6 rising to 67% at week 32.

Stone, J.H., et al., 2025. Inebilizumab for treatment of IgG4-related disease. New England Journal of Medicine 392, 1168–1177. doi:10.1056/NEJMoa2409712. pMID: 39541094. MITIGATE, n=135. Flare 10% vs 60%, HR 0.13.

Subramanian, A., Tamayo, P., Mootha, V.K., Mukherjee, S., Ebert, B.L., Gillette, M.A., Paulovich, A., Pomeroy, S.L., Golub, T.R., Lander, E.S., Mesirov, J.P., 2005. Gene set enrichment analysis: A knowledge-based approach for interpreting genome-wide expression profiles. Proceedings of the National Academy of Sciences 102, 15545–15550. doi:10.1073/pnas.0506580102.

Szklarczyk, D., Kirsch, R., Koutrouli, M., Nastou, K., Mehryary, F., Hachilif, R., Gable, A.L., Fang, T., Doncheva, N.T., Pyysalo, S., Bork, P., Jensen, L.J., von Mering, C., 2023. The STRING database in 2023: Protein–protein association networks and functional enrichment analyses for any sequenced genome of interest. Nucleic Acids Research 51, D638–D646. doi:10.1093/nar/gkac1000.

Triggianese, P., Senter, R., Perego, F., et al., 2024. Rare connective tissue diseases in patients with C1-inhibitor deficiency hereditary angioedema: First evidence on prevalence and distribution from a large Italian cohort study. Frontiers in Immunology 15, 1461407. doi:10.3389/fimmu.2024.1461407. pMID: 39493762. 4 of 18 C1INH-HAE patients with a rare connective tissue disease had primary Sjögren’s.

Verstappen, G.M., Pringle, S., Bootsma, H., Kroese, F.G.M., 2021. Epithelial–immune cell interplay in primary Sjögren syndrome salivary gland pathogenesis. Nature Reviews Rheumatology 17, 333–348. doi:10.1038/s41584-021-00605-2.

Wishart, D.S., Feunang, Y.D., Guo, A.C., Lo, E.J., Marcu, A., Grant, J.R., Sajed, T., Johnson, D., Li, C., Sayeeda, Z., Assempour, N., Iynkkaran, I., Liu, Y., Maciejewski, A., Gale, N., Wilson, A., Chin, L., Cummings, R., Le, D., Pon, A., Knox, C., Wilson, M., 2018. DrugBank 5.0: A major update to the DrugBank database for 2018. Nucleic Acids Research 46, D1074–D1082. doi:10.1093/nar/gkx1037.

Xiang, N., Xu, H., Zhou, Z., Wang, J., Cai, P., Wang, L., Tan, Z., Zhou, Y., Zhang, T., Zhou, J., Liu, K., Luo, S., Fang, M., Wang, G., Chen, Z., Guo, C., Li, X., 2023. Single-cell transcriptome profiling reveals immune and stromal cell heterogeneity in primary Sjögren’s syndrome. iScience 26, 107943. doi:10.1016/j.isci.2023.107943.

Xu, D., et al., 2024. Telitacicept in patients with primary Sjögren’s syndrome: A randomised, double-blind, placebo-controlled, phase 2 trial. Rheumatology (Oxford) 63, 698. doi:10.1093/rheumatology/kead265. pMID: 37399108. Chinese NMPA approval granted 8 June 2026 – the first regulatory approval in this indication.

Yang, M., et al., 2026. Comparative efficacy of targeted therapies in Sjögren’s disease: A systematic review and network meta-analysis. Zeitschrift für Rheumatologie doi:10.1007/s00393-026-01824-2. pMID: 42113283. 27 RCTs, 2261 patients. No agent outperformed placebo on ESSPRI.

Yu, M., et al., 2014. EphB2 contributes to human naive B-cell activation and is regulated by miR-185. The FASEB Journal 28, 3609–3617. doi:10.1096/fj.13-247759. pMID: 24803541. EphB2 knockdown reduced B-cell proliferation (-22%), TNF-alpha (-40%) and IgG production.

Zdrazil, B., Felix, E., Hunter, F., Manners, E.J., Blackshaw, J., Corbett, S., de Veij, M., Ioannidis, H., Lopez, D.M., Mosquera, J.F., Magarinos, M.P., Bosc, N., Arcila, R., Kizilören, T., Gaulton, A., Bento, A.P., Adasme, M.F., Monecke, P., Landrum, G.A., Leach, A.R., 2024. The ChEMBL Database in 2023: A drug discovery platform spanning multiple bioactivity data types and time periods. Nucleic Acids Research 52, D1180–D1192. doi:10.1093/nar/gkad1004.

Zeng, Q., Wen, J., Zheng, L., Zeng, W., Chen, S., Zhao, C., 2022. Identification of immune-related diagnostic markers in primary Sjögren’s syndrome based on bioinformatics analysis. Annals of Translational Medicine 10, 513. doi:10.21037/atm-22-1494. pMID: 35571446. CYSLTR1 is one of 8 genes overexpressed in pSS, confirmed by qRT-PCR; CYSLTR1 and TNFSF13B (BAFF) had the highest ROC sensitivity and specificity. Independent, expression-based convergent validation of the present network prediction.

Zhang, Y., Zhao, L., Sun, Y., 2024. Using single-sample networks to identify the contrasting patterns of gene interactions and reveal the radiation dose-dependent effects in multiple tissues of spaceflight mice. npj Microgravity 10, 45. doi:10.1038/s41526-024-00383-7.

Zhao, T., Zhang, R., Li, Z., Qin, D., Wang, X., 2024. Novel and potential future therapeutic options in Sjögren’s syndrome. Heliyon 10. doi:10.1016/j.heliyon.2024.e38803.

